# Carbapenem-resistant *Pseudomonas aeruginosa* bloodstream infection: *in vitro* synergy, virulence and clinical outcome

**DOI:** 10.1101/480236

**Authors:** Jessica Fernandes Ramos, Gleice Cristina Leite, Camila Rizek, Flavia Rossi, Thais Guimarães, Sabri Saeed Sanabani, Anna Sara Levin, Vanderson Rocha, Silvia Figueiredo Costa

**Affiliations:** Department of Infectious Diseases of Faculdade de Medicina, University of Sao Paulo, Brazil.; Laboratory of Medical Investigation – LIM 54 - Medical Tropical Institute, University of São Paulo, Brazil.; Laboratory of Clinical Microbiology of Hospital das Clínicas, Faculdade de Medicina, University of Sao Paulo, Brazil.; Infection Control Committee of Hospital das Clínicas, Faculdade de Medicina, University of Sao Paulo, Brazil.; Department of Haematology, Hemotherapy and Cellular Therapy of Faculdade de Medicina, University of Sao Paulo, Brazil.; Laboratory of Medical Investigation 56 – LIM 56, Medical Tropical Institute, University of São Paulo, Brazil.; Haematology Department, NHS BT, Oxford University, Oxford, UK.

**Keywords:** Bloodstream infection, Hematopoietic stem cell transplantation, multidrug-resistance, *in vitro* synergy

## Abstract

In the current study carbapanem-resistant *P. aeruginosa* bloodstream infection (CRPA-BSI) was associated with high 14-day mortality. Although, highly resistant to meropenem, half of the isolates achieved *in vitro* synergy with the combination of meropenem with colistin, and the patients that received this combination therapy showed a tendency towards lower mortality. Our strains did not carry *exo*U gene, on the other hand, patients with CRPA-BSI caused by strains harboring gene *las*B evolved more frequent to death.

## Introduction

Carbapenem-resistant *Pseudomonas aeruginosa* bloodstream infection (CRPA-BSI) has been associated with high morbidity and mortality in hematopoietic stem cell transplantation (HSCT) patients. Nowadays, it is a challenge with limited therapeutic options (1–2). Thus, *in vitro* synergy studies are important to define combination therapy that can be useful to treat these infections. To date, few studies evaluated antimicrobial synergy against CRPA (3 5) and the role of virulence on the high mortality associated with CRPA-BSI is controversial as well (2, 3).

Therefore, we aimed to describe the clinical data of 30 patients with CRPA-BSI from January 2012 to December 2014, at the Hospital das Clínicas of University of Sao Paulo, Brazil. *In vitro* synergy, pulsed field gel electrophoresis (PFGE) and PCR for carbapenemases (SPM; KPC, VIM, NDM) and virulence genes (*lasA* – protease *lasB* – elastase; *exoS* – exoenzyme S; *toxA* – exotoxin A; *phzM* – phenazine-specific methyltransferase) of the 30 strains, including those identified during the outbreak that occurred in our hospital in 2012,^2^ were performed as described elsewhere. Based on PFGE profile, whole genome sequencing (WGS) of five strains was carried out by Nextera XT, using Illumina MiSeq technology. *De novo* assembly of reads was performed using VelvetOptimiser v.2.2.5 (Victorian Bioinformatics Consortium, Australia) and contigs were ordered by Abacas v.1.3.1. The annotation of genome was performed by Prokka v.1:11. *P. aeruginosa* PAO1 was used as reference. The antibiotic resistance and virulence genes analysis was performed using the following tools: Resfinder (https://cge.cbs.dtu.dk/services/ResFinder/), Blast2Seq (https://blast.ncbi.nlm.nih.gov/Blast.cgi) and MAFFT (http://mafft.cbrc.jp/alignment/server/). Subsequently, the presence of the genes were confirmed by manual curation of CDS and by Artemis program The Multilocus Sequence Typing (MLST) analysis was performed by MLSTfinder tool (https://cge.cbs.dtu.dk/services/MLST/) and analyzed by software eBURST (http://eburst.mlst.net/).

BSI definition was based on Centres for Diseases Control and Prevention (CDC) criteria (6) and Pitt bacteraemia score was calculated (7). All patients received levofloxacin as prophylaxis during neutropenia. Initial antibiotic therapy was considered appropriate when colistin (COL) was administered within 24 hours after obtain blood culture. All patients received 2g of meropenem (MERO) tid, prolonged infusions (3 hours) adjusted for kidney function when indicated. Amikacin (AMK) was added to therapy with colistin and meropenem whenever CRPA was susceptible.

Minimum inhibitory concentrations (MIC) of COL (USP Reference Standard, Rockville, MD, USA), AMK (Sigma-Aldrich, St Louis, MO, USA), and MERO (Astra Zeneca, Cotia, SP, Brazil) were determined using the broth microdilution method according to the Clinical and Laboratory Standards Institute (8). *Time-kill* assay was performed in duplicate with drugs alone and combined at 1x MIC and 0.5x MIC as previously reported (9–10). Aliquots were removed at time 0 and 2, 4, 6, and 24 hours and serially diluted in 0.85% sodium chloride solution. Diluted samples of 0.01 mL were plated on Müeller-Hinton agar, and the colonies were counted (log_10_ cfu/mL) after 20 hours of incubation at 37°C. Synergy was interpreted as a ≥ 2 log_10_ decrease in colony count with the combination compared to the most active single agent; was considered antagonistic for a ≥ 2 log_10_ increase in cfu/mL, and indifferent for a < 2 log_10_ increase or decrease in count (9–10). Bivariate Cox regressions were used to evaluated 14-day mortality using SPSS software version 21 (SPSS Inc., USA), value of p less than 5% was considered significant.

The strains presented MERO MIC90 > 512µg/mL; two thirds were also resistant to AMK (MIC 2-512 µg/mL) and all were susceptible to COL (Table 1). The combination of COL plus MERO achieved synergy in 57% of isolates, COL plus AMK in 2/30 isolates (6.6%), and of MERO plus AMK in 33% (10/30), only in isolates susceptible to AMK (table 2). Although, highly resistant to meropenem, half of the isolates achieved *in vitro* synergy with combination of meropenem with colistin, and the group that received this combination showed a tendency towards lower mortality (6 of 7 patients that survived received meropenem plus colistin) (p=0.06). Zusman et al (2013) performed a meta-analysis and found similar results for *in vitro* combination of carbapenem and polymixin B or colistin in 59% of 43 CRPA isolates (4).

**Table 1.** Microbiological characteristics of thirty isolates of carbapenem-resistant *P. aeruginosa* of bloodstream infection.

| Isolates | PFGE<br>carbapenemase | MIC Results (mg/ml) |  |  |  |  |  | In vitro synergy by time-kill |  |  |
| --- | --- | --- | --- | --- | --- | --- | --- | --- | --- | --- |
|  |  | AMK |  | COL |  | MERO |  | COL+MERO | COL+AMK | MERO+AMK |
| 1 | D<br>SPM/ KPC | 512 | R | 0,5 | S | 256 | R | Yes | - | RG 24h |
| 2 | B<br>SPM/ KPC | 4 | S | 0,5 | S | 512 | R | Yes | RG 6h | Yes |
| 3 | B<br>SMP/ KPC | 4 | S | 0,5 | S | 256 | R | RG 24h | RG 6h | Yes |
| 4 | E<br>SPM/ KPC | 2 | S | 0,5 | S | 16 | R | Yes | RG 6h | Yes |
| 5 | B<br>SPM/ KPC | 4 | S | 1 | S | 512 | R | RG 24h | RG 6h | Yes |
| 6 | D<br>SMP | 4 | S | 1 | S | 512 | R | RG 24h | RG 24h | Yes |
| 7 | A<br>SPM | 4 | S | 1 | S | 512 | R | Yes | - | Yes |
| 8 | A<br>SPM | 8 | S | 0,5 | S | 512 | R | Yes | RG 24h | Yes |
| 9 | A<br>SPM | 512 | R | 0,5 | S | 256 | R | RG 24h | - | RG 24h |
| 10 | A<br>SPM | 512 | R | 1 | S | 256 | R | RG 24h | - | RG 24h |
| 11 | A<br>SPM | 512 | R | 1 | S | 512 | R | RG 24h | - | RG 24h |
| 12 | A<br>SPM | 512 | R | 1 | S | 256 | R | RG 24h | - | RG 24h |
| 13 | C<br>SMP/ KPC | 512 | R | 0,5 | S | 256 | R | RG 24h | - | RG 24h |
| 14 | D | 512 | R | 0,5 | S | 256 | R | Yes | - | RG 24h |
| 15 | D | 512 | R | 1 | S | 512 | R | RG 24h | - | RG 24h |
| 16 | D<br>SPM | 512 | R | 0,5 | S | 256 | R | RG 24h | - | RG 24h |
| 17 | D<br>SPM | 512 | R | 0,5 | S | >256 | R | RG 24h | - | RG 24h |
| 18 | A<br>SPM | 512 | R | 1 | S | 256 | R | Yes | - | RG 24h |
| 19 | A<br>SPM | 512 | R | 1 | S | 512 | R | RG 24h | - | RG 24h |
| 20 | A<br>SPM | 512 | R | 1 | S | 256 | R | Yes | - | RG 24h |
| 21 | A<br>SPM | 512 | R | 0,5 | S | 256 | R | RG 24h | - | RG 24h |
| 22 | A<br>SPM | 8 | S | 0,5 | S | 512 | R | Yes | Yes | Yes |
| 23 | A<br>SPM | 4 | S | 0,5 | S | 512 | R | Yes | Yes | Yes |
| 24 | A | 256 | R | 0,5 | S | 256 | R | Yes | - | RG 24h |
| 25 | A<br>SPM | 256 | R | 0,5 | S | 256 | R | Yes | - | RG 24h |
| 26 | A<br>SPM | 8 | S | 0,5 | S | 512 | R | Yes | RG 24h | Yes |
| 27 | B | 512 | R | 4 | I | 512 | R | Yes | RG 24h | RG 24h |
| 28 | B<br>SPM | 512 | R | 4 | I | 512 | R | Yes | RG 24h | RG 24h |
| 29 | B | 512 | R | 4 | I | 512 | R | Yes | RG 24h | RG 24h |
| 30 | B | 512 | R | 4 | I | 512 | R | Yes | RG 24h | RG 24h |

MIC: Minimum inhibitory concentration, S: susceptible, R: resistant, I: intermediate, COL: colistin, MERO: meropenem, AMK: amikacin, RG: bacterial re-grow, (-) no synergyism

**Table 2.**
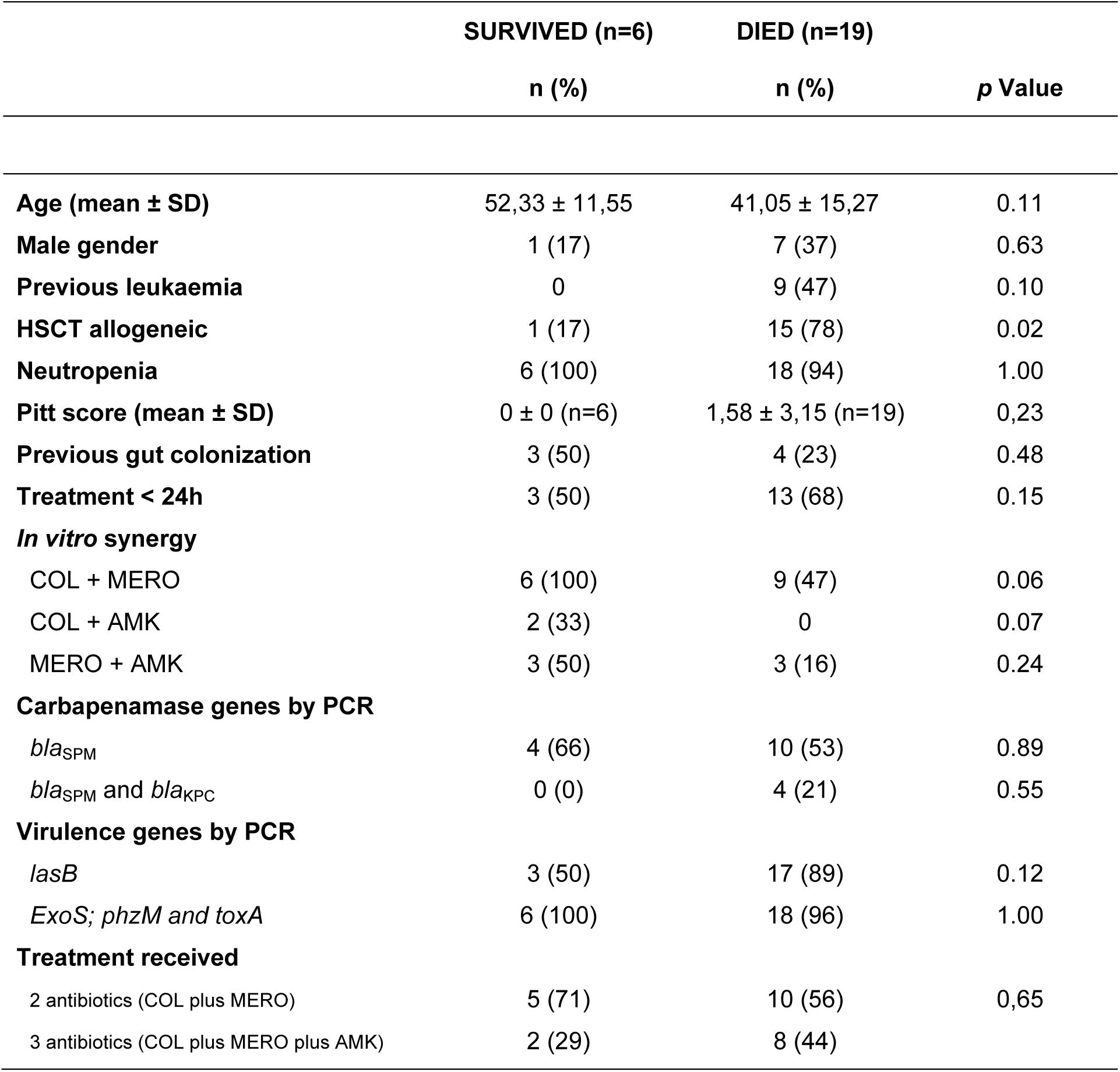
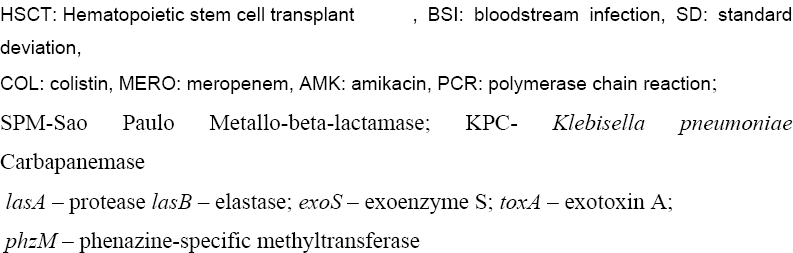
Comparison of clinical and laboratorial features of 25 patients submitted to HSCT with carbapenem-resistant *P. aeruginosa* BSI appropriated treated regarding outcome – São Paulo, Brazil.

|  | **SURVIVED (n=6)** | **DIED (n=19)** |  |
| --- | --- | --- | --- |
|  | **n (%)** | **n (%)** | **p Value** |
| **Age (mean ± SD)** | 52,33 ± 11,55 | 41,05 ± 15,27 | 0.11 |
| **Male gender** | 1 (17) | 7 (37) | 0.63 |
| **Previous leukaemia** | 0 | 9 (47) | 0.10 |
| **HSCT allogeneic** | 1 (17) | 15 (78) | 0.02 |
| **Neutropenia** | 6 (100) | 18 (94) | 1.00 |
| **Pitt score (mean ± SD)** | 0 ± 0 (n=6) | 1,58 ± 3,15 (n=19) | 0,23 |
| **Previous gut colonization** | 3 (50) | 4 (23) | 0.48 |
| **Treatment < 24h** | 3 (50) | 13 (68) | 0.15 |
| ***In vitro* synergy** |  |  |  |
| COL + MERO | 6 (100) | 9 (47) | 0.06 |
| COL + AMK | 2 (33) | 0 | 0.07 |
| MERO + AMK | 3 (50) | 3 (16) | 0.24 |
| **Carbapenamase genes by PCR** |  |  |  |
| *bla*<sub>SPM</sub> | 4 (66) | 10 (53) | 0.89 |
| *bla*<sub>SPM</sub> and *bla*<sub>KPC</sub> | 0 (0) | 4 (21) | 0.55 |
| **Virulence genes by PCR** |  |  |  |
| *lasB* | 3 (50) | 17 (89) | 0.12 |
| *ExoS*; *phzM* and *toxA* | 6 (100) | 18 (96) | 1.00 |
| **Treatment received** |  |  |  |
| 2 antibiotics (COL plus MERO) | 5 (71) | 10 (56) | 0,65 |
| 3 antibiotics (COL plus MERO plus AMK) | 2 (29) | 8 (44) |  |

HSCT: Hematopoietic stem cell transplant , BSI: bloodstream infection, SD: standard deviation, COL: colistin, MERO: meropenem, AMK: amikacin, PCR: polymerase chain reaction; SPM-Sao Paulo Metallo-beta-lactamase; KPC- *Klebisella pneumoniae* Carbapanemase *lasA* – protease *lasB* – elastase; *exoS* – exoenzyme S; *toxA* – exotoxin A; *phzM* – phenazine-specific methyltransferase

Clinical and demographic data are shown in table 2. Mortality was higher among allogeneic HSCT recipients compared to autologous recipients (79% vs. 17% p = 0,012). Patients treated with two or three drugs did not present a statistically significant difference in 14-day death (29 % x44% p = 0,65) (table 2).

Five clones named as A, B, C, D and E harboured SPM and belonged to ST277 were found by PFGE during the study period (table 1). ST277 is common in Brazil and has been described in several outbreaks including patients who travelled to the country (13–16). Interestingly, the six strains that co-harbored KPC were distributed in four clones, except the predominant clone named as clone A. We did not find *exo*U in our strains, on the other hand, patients with CRPA BSI caused by strains harboring gene *las*B evolved more frequent to death (88.9% vs. 57.1%). This virulence factor is associated with bacterial enzyme production of elastase that degrades immunoglobulin and complement factor (12). In Brazil, the gene *exo*U was described in CRPA belonging to ST2237 in burn patients instead of ST277 (17).

The most frequent carbapenemase identified was *bla*_SPM_, and six isolates co-harboured *bla*_KPC_ as shown in table 1. The five clones sequenced by illumina were assigned as ST277 and harbored the Tn4371 (Table 3). In all the strains, mutations on outer membrane protein OprD (T103S, K115T, V118P, F170L) OprM (A261T), OprJ (D68G, M69V) were found, as well as the *mexT* gene that possessed a mutation resulting in a frameshift. The 16S rRNA methyltransferase gene rmtD1, which confers high-level resistance to all aminoglycosides and has been associated to ST277 was present only in clones I, IV and V. Clone III did not harbor *fos*A, it carried *exo*S and *tox*A of Type Three Secretion System (TTSS) and lacked the *exo*Y gene. Genbank access number are QHLR00000000, QHLU00000000, QHLW00000000, QJPG00000000, and QJPA00000000.

**Table 3.**
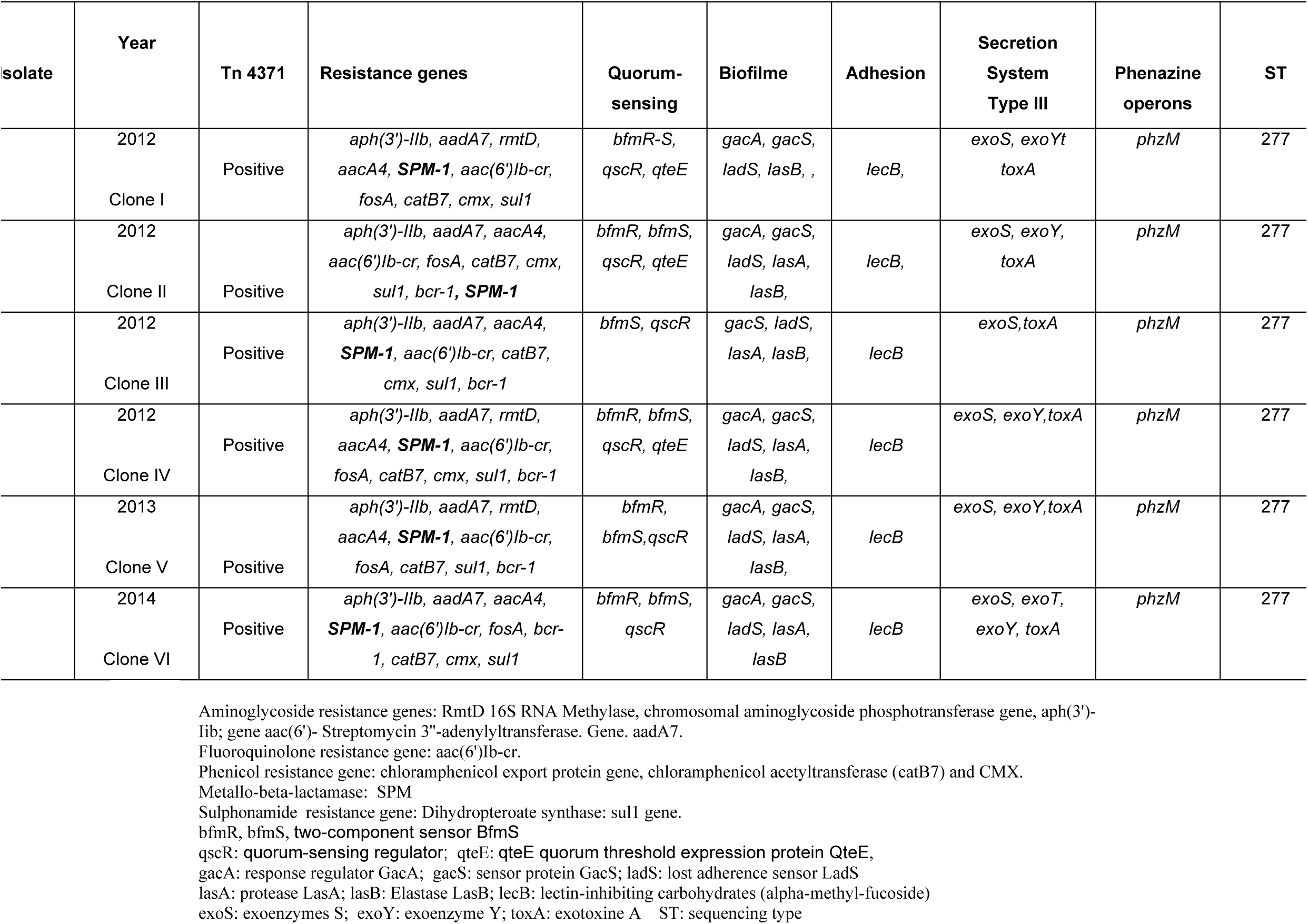
Resistance and virulence genes by whole genome sequence of main clones of carbapenem-resistant *P. aeruginosa* casing bloodstream infection in HSCT patients.

Although, the study was underpowered to recommended combination therapy with more than two drugs, it highlights the importance to test *in vitro* synergy to treat infections caused by CRPA.

## Ethical statement

The ethical committee of Hospital das Clinicas and CONEP (National Ethics Commission), Brazil, gave ethical approval.

## Author contribution statements

JR collected and analysed data and written the article, GCL performed time kill and PFGE, CR performed PCR, RR analysed the whole genome sequence, SS performed de whole genome sequence, FR provides the strains, TG provides clinical data, ASL analysed data and reviewed the article, VR provide clinical data, SFC design the study, analysed data and written the article.

## REFERENCES

1. Andria N, Henig O, Kotler O, Domchenko A, Oren I, Zuckerman T, Ofran Y, Fraser D, Paul M. 2015. Mortality burden related to infection with carbapenem-resistant Gram-negative bacteria among hematological cancer patients: a retrospective cohort study. J Antimicrob Chemother; 70: 3146–3153.

2. Chaves L, Tomich LM, Salomão M, Leite GC, Ramos J, Martins RR, Rizek C, Neves P, Batista MV, Amigo U, Guimaraes T, Levin AS, Costa SF et al. 2017. High mortality of bloodstream infection outbreak caused by carbapenem-resistant P. aeruginosa producing SPM-1 in a bone marrow transplant unit. Journal of Medical Microbiology; 66: 1722–1729.

3. Peña C, Cabot G, Gómez-Zorrilla S, Zamorano L, Ocampo-Sosa A, Murillas J, Almirante B, Pomar V, Aguilar M, Granados A, Calbo E, Rodríguez-Baño J, Rodríguez-López F, Tubau F, Martínez-Martínez L, Oliver A; 2015. Spanish Network for Research in Infectious Diseases (REIPI). Influence of virulence genotype and resistance profile in the mortality of Pseudomonas aeruginosa bloodstream infections. Clin Infect Dis 4:539–548.

4. Zusman O, Avni T, Leibovici L, Adler A, Friberg L, Stergiopoulou T, Carmeli Y, Paul M. et al. 2013. Systematic Review and Meta-Analysis of *In Vitro* Synergy of Polymyxins and Carbapenems. Antimicrobial Agents and Chemotherapy; 57 (10): 5104–5111.

5. He W, Kaniga K, Lynch AS, Flamm RK, Davies TA. 2012. In vitro Etest synergy of doripenem with amikacin, colistin, and levofloxacin against Pseudomonas aeruginosa with defined carbapenem resistance mechanisms as determined by the Etest method. Diagnostic Microbiology and Infectious Disease; 74: 417–419.

6. Horan TC, Andrus M, Dudeck MA 2008. CDC/NHSN surveillance definition of health care–associated infection and criteria for specific types of infections in the acute care setting. Am J Infect Control; 36: 309–332.

7. Rhee JY, Kwon KT, Ki HK, Shin SY, Jung DS, Chung DR, Ha BC, Peck KR, Song JH. 2009. Systems for prediction of mortality in patients with intensive care unit acquired sepsis: a comparison of the Pitt bacteremia score and the Acute Physiology and chronic health evaluation II scoring systems. Shock; 31 (2): 146–150.

8. Clinical and Laboratory Standard Institute (2013) Performance standards for antimicrobial susceptibility testing. Nineteenth informational supplement. CLSI document M100 S19. CLSI, Wayne, PA, USA.

9. Petersen PJ, Labthavikul P, Jones CH, Bradford PA. 2006. In vitro antibacterial activities of tigecycline in combination with other antimicrobial agents determined by checkerboard and time-kill kinetic analysis. J Antimicrob Chemother; 57(3): 573–576.

10. Pillai SK, Moellering RC, Eliopoulos GM. 2005. In: Antimicrobial combinations. Antibiotics in laboratory medicine. Lorian V, Ed. 5th ed. Philadelphia, PA: Lippincott Williams and Wilkins.

11. Hu Y, Li L, Li W, Xu H, He P, Yan X, Dai Hl. 2013. Combination antibiotic therapy versus monotherapy for *Pseudomonas aeruginosa* bacteraemia: a meta-analysis of retrospective and prospective studies. Int J Antimicrob Agents 42: 492–496.

12. Pereira SG, Rosa AC, Cardoso O. 2015. Virulence factors as predictive tools for drug resistance in *Pseudomonas aeruginosa*. Virulence 5: 1– 4.

13. Araujo BF, Ferreira ML, Campos PA, Royer S, Batistão DW, Dantas RC, Gonçalves IR, Faria AL, Brito CS, Yokosawa J, Gontijo-Filho PP, Ribas RM. 2016. Clinical and Molecular Epidemiology of Multidrug-Resistant P. aeruginosa Carrying aac(6’)-Ib-cr, qnrS1 and blaSPM Genes in Brazil. PLoS One 5:e0155914.

14. Nascimento AP, Ortiz MF, Martins WM, Morais GL, Fehlberg LC, Almeida LG, Ciapina LP, Gales AC, Vasconcelos AT.2016. Intraclonal Genome Stability of the Metallo-β-lactamase SPM-1-producing Pseudomonas aeruginosa ST277, an Endemic Clone Disseminated in Brazilian Hospitals. Front Microbiol 7:1946.

15. Hopkins KL, Meunier D, Findlay J, Mustafa N, Parsons H, Pike R, Wright L, Woodford N. 2016. SPM-1 metallo-β-lactamase-producing Pseudomonas aeruginosa ST277 in the UK. J Med Microbiol 7:696– 697.

16. Salabi AE, Toleman MA, Weeks J, Bruderer T, Frei R, Walsh TR. 2010. First report of the metallo-beta-lactamase SPM-1 in Europe. Antimicrob Agents Chemother 1:582.

17. de Almeida Silva KCF, Calomino MA, Deutsch G, de Castilho SR, de Paula GR, Esper LMR, Teixeira LA. 2017. Molecular characterization of multidrug-resistant (MDR)Pseudomonas aeruginosa isolated in a burn center. Burns 1:137–143.

